# A plant immune receptor mediates tritrophic interactions by linking caterpillar detection to predator recruitment

**DOI:** 10.1101/2025.07.29.667524

**Authors:** Natalia Guayazán Palacios, Patrick Grof-Tisza, Brian Behnken, Carla Marques Arce, Di Wu, Antonio F. Chaparro, Eric A. Schmelz, Ted C.J. Turlings, Betty Benrey, Adam D. Steinbrenner

**Affiliations:** Department of Biology, University of Washington. Seattle, WA, USA; Laboratory of Evolutionary Entomology, Institute of Biology, University of Neuchâtel, Neuchâtel, 2000 Switzerland; Department of Natural Sciences, Converse University, Spartanburg, 29032 SC, USA; Department of Plant & Microbial Biology, University of California Berkeley. Berkeley, CA, USA; Section of Cell and Developmental Biology, University of California, San Diego, La Jolla, CA, USA; State Key Laboratory of Crop Stress Adaptation and Improvement, State Key Laboratory of Cotton Bio-breeding and Integrated Utilization, School of Life Sciences, Henan University, Kaifeng, 475004, China; Laboratory of Fundamental and Applied Research in Chemical Ecology, University of Neuchȃtel, Neuchȃtel 2000, Switzerland; Department of Entomology, The Pennsylvania State University, University Park, Pennsylvania, USA

## Abstract

Plants deploy direct and indirect defenses in response to insect herbivory. The specific antiherbivore responses involve cell surface immune receptors that recognize herbivore associated molecular patterns (HAMPs), yet the ecological relevance of this molecular interplay remains unexplored. We demonstrate that the Inceptin Receptor (INR) encoded by common bean, upon recognition of inceptin, a HAMP in caterpillar oral secretions, orchestrates a tritrophic interaction. Laboratory and field experimentation in Mexico using a naturally occurring inceptin-insensitive mutant and its near-isogenic wild type equivalent, revealed that inceptin recognition by INR activates an herbivore-specific immune pathway, and triggers the emission of a volatile blend that recruits predatory wasps. These findings provide a novel molecular-to-ecological link, revealing how an immune receptor mediates ecologically relevant plant–insect–predator interactions in nature.

## Main

Plants defend themselves against antagonists through complex immune responses that are initiated when cell surface receptors recognize conserved molecular patterns^1–7^. These include herbivore-associated molecular patterns (HAMPs) present in the oral secretions and eggs of herbivorous insects and that are deposited during feeding and oviposition^6–8^. The presence of HAMPs can activate direct defenses in the form of chemical and physical changes that directly reduce herbivore performance^9,10^, as well as indirect defenses like the emission of volatiles that attract natural enemies of the herbivores^11,12^. Several defense-eliciting HAMPs have been identified from insect oral secretions, including β-glucosidases^13^, fatty acid-amino acid conjugates^14^, caeliferins^15^, mucin-like proteins^16^, vitellogins^17^ and inceptins^18^. Yet, how plants perceive these HAMPs has long remained unclear. Recently the Inceptin Receptor (INR), a leucine-rich repeat receptor-like protein (LRR-RLP) found in various Phaseoloid legumes, was identified as the receptor of inceptins, including the 11 amino acid peptide In11^19,20^. In11 is a fragment of the chloroplast ATP synthase γ-subunit that is generated upon ingestion of leaf tissue by caterpillars and perceived at picomolar concentrations by the leaves of legumes^18^. While biochemical and molecular studies have elucidated INR’s role in initiating immune signaling and transcriptional reprogramming^19,21,22^, the broader ecological implications of inceptin perception remain unknown. Here, we exploit natural variation in ln11 perception to demonstrate in laboratory and field experiments that INR mediates direct defense activation as well as the emission of specific volatile blends that recruit natural enemies in a realistic agricultural setting. Our findings establish a direct link between the molecular recognition of herbivores and ecologically relevant tritrophic outcomes, providing insight into the ecological processes that shape the evolution of plant immune systems.

### Natural variation in In11-induced defense is associated with a deletion in the INR gene

The lack of efficient gene transformation or silencing methods in common bean has limited reverse genetic study of INR’s role in induced defenses. Instead, we aimed to identify naturally occurring In11-insensitive common bean varieties. We screened a panel of Mesoamerican landraces for characteristic induced ethylene gas accumulation in response to wounding (w + H_2_O) and application of In11 (w + In11)^18^. While 89 varieties had In11/H_2_O induced ethylene ratio of 1.2 or higher, two insensitive landraces, Honduran landrace W6 13807 and Argentinian landrace W6 17491, did not produce induced ethylene in response to In11 (In11/H_2_O ≤ 1) (Extended data Table 1). We focused on Honduran landrace W6 13807 and, to facilitate crossing and generation of germplasm, we also identified a sexually compatible, closely related landrace PI 311785 with standard In11-induced ethylene response (Fig. 1a, Extended data Fig. 1a).

**Figure 1.**
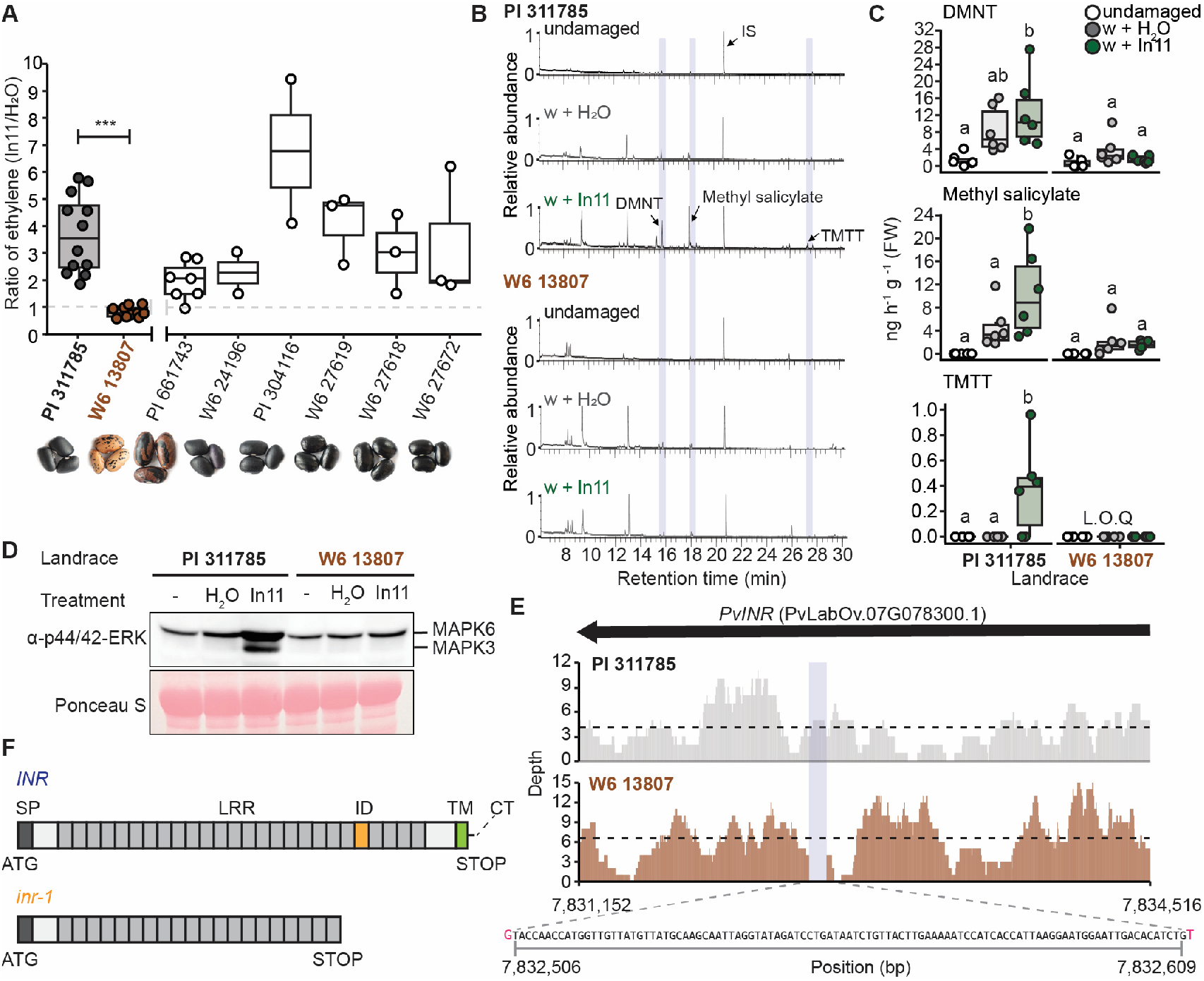
*inr-1* is a naturally occurring deletion allele among Honduran bean landraces. **(A)** Ratio of ethylene gas induced by wounding (w) + 1 µM In11 treatment relative to mock treatment (w + H_2_O) for select common bean (*Phaseolus vulgaris*) landraces. Seed coat diversity among landraces is shown. Boxplots represent the 1^st^ and 3^rd^ quantile and the median. Each dot represents an independent plant. Comparing varieties PI 311785 and W6 13807 used for crossing and introgression, significant differences in the mean were determined by a two-sided t-test (n = 10-12, α= 0.05), ***, p<0.001. (**B)** Representative GC-MS chromatograms of volatiles collected from bean varieties PI 311785 and W6 13807 after the plants had either been left unharmed (undamaged), wounded and treated with water (w + H_2_O), or wounded and treated with 1 µM ln11 (w + In11). (**C)** Quantified concentrations of individual volatile organic compounds. Different letters indicate significant differences within treatments of each variety. Linear model [family, Gaussian] followed by pairwise comparisons of least squares means. L.O.Q = Limit of quantification. (**D)** MAPK 3 and 6 phosphorylation measured 15’ after spraying H_2_O or 1 µM In11 on indicated landraces. Protein loading is shown by Ponceau stain. **(E)** Read depth coverage of re-sequenced PI 311785 and W6 13807 landraces mapped to the *PvINR* locus in the Mesoamerican (Guatemalan) *Phaseolus vulgaris* Labor Ovalle v1.1 genome. The average read depth is shown as dashed lines. The sequence and position of the 103-base pair (bp) deletion is also indicated with a vertical bar. The nucleotides flanking the deletion are shown in magenta. (**F)** Domain architecture of INR receptor and *inr-1* deletion variant with resulting frameshift mutation. SP: signal peptide, ID: Island domain, TM: Transmembrane domain, CT: Cytoplasmic tail.

The emission of specific blends of herbivore-induced volatiles (HIPVs) to attract natural enemies has been recognized as an important form of indirect plant defense^23–26^, as HIPVs induced by live herbivores or artificial hormone treatment can reduce oviposition and increase predation rates in the field^27–29^. The perception of In11 in a related species (cowpea) results in induced expression of biosynthetic terpene synthase (TPS) genes and the accumulation of specific volatile blends that include the homoterpenes (E)-4,8-dimethyl-1,3,7-nonatriene (DMNT) and (3E,7E)-4,8,12-trimethyl-1,3,7,11-tridecatetraene (TMTT)^18,21^, important signaling compounds known to mediate indirect plant defenses^30–32^. We hypothesized that HIPV emission is impaired in In11-insensitive common bean plants. Indeed, we found that the responsive variety PI 311785 emitted a distinct volatile profile in response to wounding plus In11 compared to wounding alone, whereas W6 13807 responded with a much-reduced emission that was consistent with lack of In11 recognition (Extended data Fig. 1b and c). Specifically, methyl salicylate (MeSA) was significantly higher in PI 311785 but not W6 13807 in response to In11, while the less abundant TMTT was uniquely present in In11-treated PI 311785, and DMNT showed a similar trend (Fig. 1b and Fig. 1c). These volatiles are known to serve as foraging cues for natural enemies, contributing to indirect plant defenses^28,30,33^.

We additionally measured MAPK3/6 phosphorylation, a canonical marker of plant pattern triggered immunity (PTI)^34,35^, in response to spray application of In11. We found that MAPK3 and MAPK6 orthologs were phosphorylated in PI 311785 but not in W6 13807 15 minutes after elicitation with In11 (Fig. 1d). Together, these results suggested a lack of activation in the In11-specific signaling cascade upstream of MAPK phosphorylation that might be required for HIPV emission in W6 13807.

To identify the potential genetic basis of In11 insensitivity, we generated short read re-sequencing data for W6 13807. Mapping to the Mesoamerican reference genome *P. vulgaris* cv. Labor Ovalle revealed a candidate 103 base pair (bp) deletion in W6 13807 without read support in the LRR domain of the Inceptin Receptor (INR) (Fig. 1e). The resulting protein product is expected to be truncated at the last LRR before the island domain (Fig. 1f), indicating that attachment to the plasma membrane and ligand-induced interaction with co-receptors is impaired^36,37^. Furthermore, F2 populations from the W6 13807 x PI 311785 cross showed segregation of inducible ethylene accumulation, indicating that sensitivity to In11 is conferred by a simply inherited locus. Genotyping of F2 individuals revealed co-segregation of the 103-bp deletion with response to In11 (Extended data Fig. 1d). Consistent with these phenotypic observations for common bean, in a previous study the presence of a functional INR homolog corresponded with In11 sensitivity across 20 diverse legume species^20^. Thus, we conclude that In11 insensitivity in common bean landrace W6 13807 is associated with a naturally occurring null allele, which we termed *inr-1*.

### Direct defenses are impaired in *inr-1* deletion lines

To directly compare the effect of the *inr-1* allele on direct and indirect defenses, we generated three sets of near-isogenic lines (NILs) through independent W6 13807 x PI 311685 crosses, followed by six recurrent backcrosses with PI 311785, and final selection of *inr-1/inr-1* and *INR/INR* genotypes (Extended data Fig. 2a). Consistent with data from parent lines and F2 individuals, the *inr-1/inr-1* NILs did not produce In11-induced ethylene (Fig. 2a) but retained the ability to recognize the bacterial peptide flg22 (Extended data Fig. 2b), indicating that insensitivity to In11 is not due to a downstream signaling component required for immune responses. Importantly, *Spodoptera exigua* larval growth rate was 72.7% higher on *inr-1/inr-1* relative to *INR/INR* sibling NILs over a 5-day feeding time course (Fig. 2b). Together, these results imply that a functional allele of *INR* is needed for In11 to induce direct defense responses that enhance herbivore resistance in common bean.

**Figure 2.**
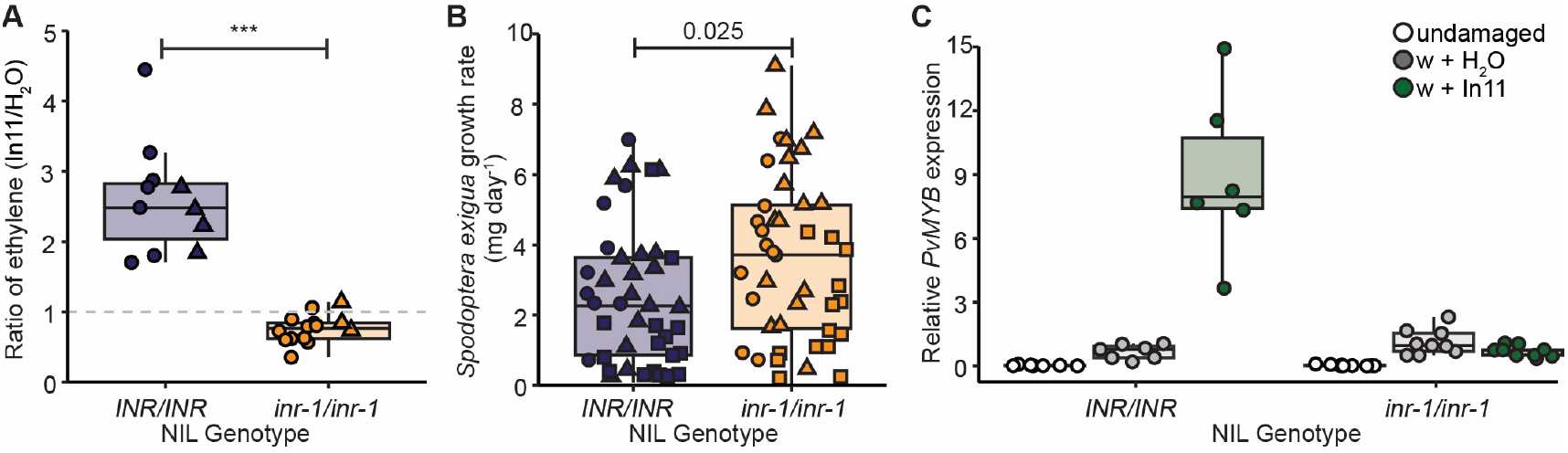
*inr-1* is associated with reduced In11-induced defense responses and increased herbivore growth in near-isogenic lines. **(A)** Ratio of ethylene gas induced by wounding (w) + 1 µM In11 treatment relative to mock treatment (w + H_2_O) for common bean near-isogenic lines (NILs) with *INR/INR* or *inr-1/inr-1* genotype. Each dot represents an independent plant (n=11-15). Boxplots represent the 1^st^ and 3^rd^ quantile and the median. Significant differences in the mean were determined by a two-sided t-test (n = 11-13, α= 0.05), ***, p<0.001. (**B)** Growth rate of 2^nd^ instar *Spodoptera exigua* larvae feeding on common bean NILs for 5 days. Each dot represents one larva feeding on an individual plant. Significant differences in the mean were determined by a two-sided t-test (n = 40-43, α= 0.05). Symbols in panels A and B indicate data from independent experiments using a mixture of NILs 3, 4 and 8 (Extended data Fig. 2c). (**C)** *PvMYB* (*Phvul*.*001G215100*) marker gene expression 1 hour after wounding and adding water or 1 µM In11. Gene expression was normalized against *PvUBQ*.

To identify downstream components of the INR-dependent immune pathway, we generated RNA sequencing data from *P. vulgaris* cv. Red Hawk 1 hour after treatment with wounding + In11 compared to mock treatment (w + H_2_O). Our analysis revealed In11-specific gene expression (Extended data Fig. 3a and Supplementary Table 1) defined by 527 genes that were up-regulated by In11 treatment relative to wounding alone (Extended data Fig. 3b). From these, 29.6% belonged to Gene Ontology (GO) categories associated with protein phosphorylation and regulation of transcription (Extended data Fig. 3c and Supplementary Table 2). We selected the transcription factor *PvMYB* as an induced marker gene and found that it was highly upregulated in *INR/INR* but not in *inr-1/inr-1* NILs 1 hour after In11 treatment (Fig. 2c), suggesting that recognition of In11 by INR results in rapid herbivore-specific transcriptional reprogramming.

### Impaired In11-induced volatile emissions in inr-1deletion lines negatively impacts predatory wasp attraction in a Mexican field

In their natural environments, plant’s emission of specific blends of HIPVs recruit natural enemies^12^ that serve as indirect defenses by predating or parasitizing herbivorous pests^38^. To determine if *INR* mediates the emission of specific HIPV blends we measured and characterized In11-induced volatiles in the common bean NILs grown under laboratory conditions.

We found that *inr-1/inr-1* NILs did not emit the typical HIPV blend that is normally induced upon In11 treatment, but rather emitted volatiles that bean plants release after wounding alone. In contrast, *INR/INR* NILs produced the characteristic blend of volatiles in response to treatment with a physiologically relevant concentration of In11 alone^18^ or total oral secretions (OS) derived from bean-fed fall armyworm (*Spodoptera frugiperda*) caterpillars (Fig. 3a). Among other compounds, DMNT, MeSA, and TMTT were predominantly detected in the headspace of In11- or OS-treated *INR/INR* NILs (Fig. 3b) and at significantly higher levels than those with the w + H_2_O treatment (Fig. 3c), thus contributing to the distinct volatile blends observed between the NIL genotypes. These results are consistent with previous reports that In11 is the primary elicitor of plant responses found in OS^18^, and indicate that INR is required for specific volatile emission in response to synthetic and natural In11 found in the caterpillar OS.

**Figure 3.**
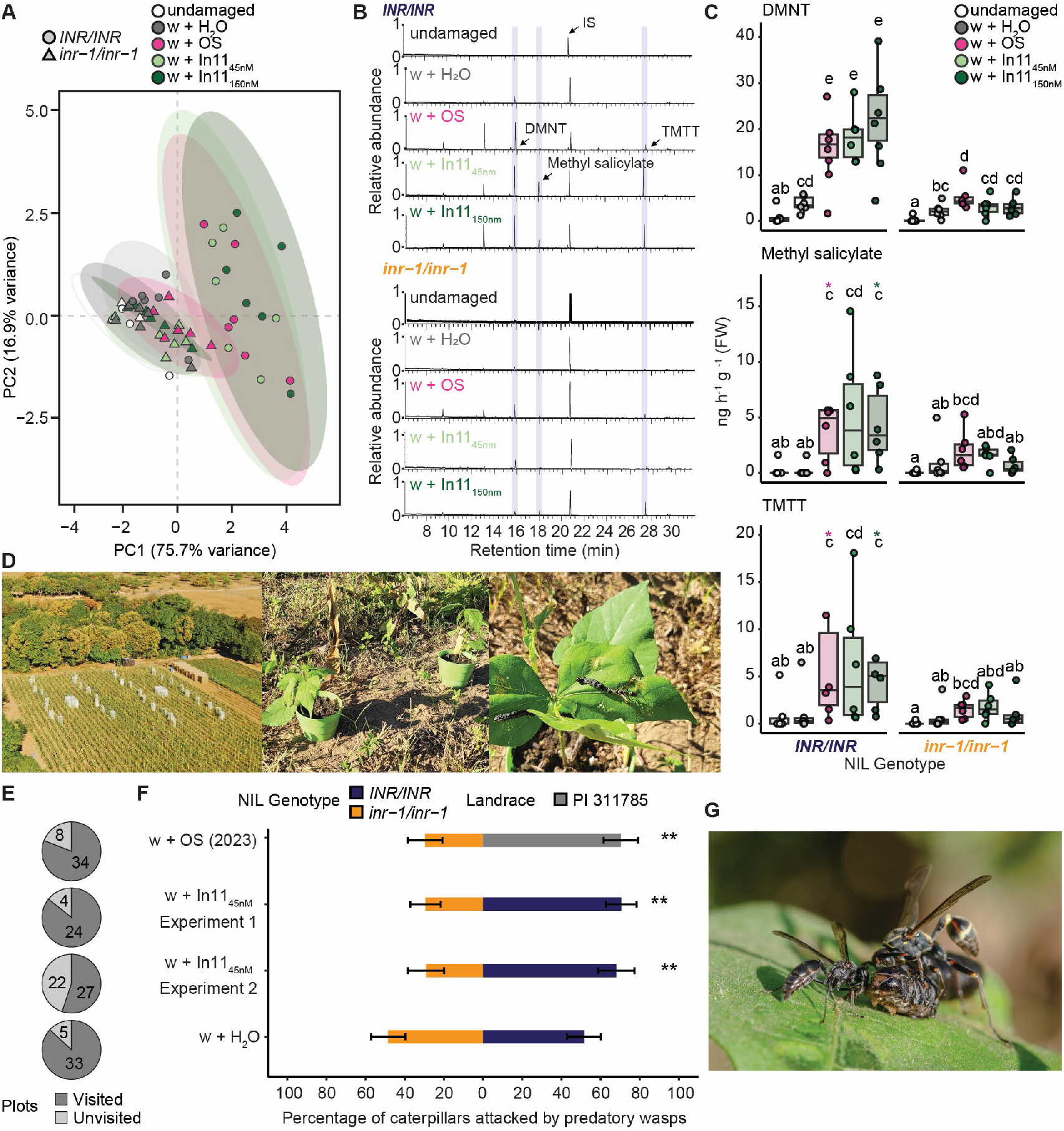
*inr-1* is associated with impaired In11-induced volatile emission and reduced attraction of predatory wasps in near-isogenic lines of common bean plants in Mexico. Panels **A–C** show results from a shared headspace volatile collection experiment comparing NILs (INR/INR and *inr-1/inr-1*) subjected to five treatments: undamaged, mechanical wounding (w + H_2_O), wounding with fall armyworm (FAW, *Spodoptera frugiperda*) oral secretions (w + OS), a physiologically relevant concentration of In11 (45 nM, w + In11_45nM_), or an excess of In11 (150 nM, w + In11_150 nM_) (n = 6). (**A**) PCA ordination plot of VOC profiles, illustrating treatment- and genotype-driven separation. Ellipses represent 95% confidence intervals. (**B)** Representative GC-MS chromatograms of headspace induced volatile organic compounds (VOCs) (**C)** Boxplots of selected VOCs, known to be important in the indirect defenses of plants, 1 hour after treatment. Letters denote significant differences from a generalized linear model (Gamma error) with pairwise least squares means comparisons (α = 0.05). Asterisks indicate extreme values not shown for clarity. (**D**) Field experimental setup in Oaxaca, Mexico, showing block layout of where wasp visits were monitored, treatment-assigned NIL pairs, and pinned sentinel FAW caterpillars. **(E)** Visitation rates to each NIL/treatment pair. (**F)** Bar plots show the mean percentage of caterpillars attacked by social wasps across treatments and two field seasons (2023, 2024). Statistical analysis was performed using a binomial generalized linear mixed model. **, p<0.01. (**G)** Representative image of a committed attack by *Polybia sp* and *Mischocyttaurus sp* on a sentinel FAW caterpillar.

Lastly, we hypothesized that a functional INR would mediate the recruitment of natural enemies to plants treated with In11 or herbivore-derived OS. To test this in nature, we adapted a sentinel-prey field assay^39^ and recorded predatory wasp visitation to pairs of common bean NILs (Fig. 3d) treated with wounding plus water, OS or a physiologically relevant concentration of In11 in an experimental agricultural field in Oaxaca, Mexico over two field seasons in 2023 and 2024. There was considerable wasp activity across both seasons as indicated by plot visitation rates ranging from 55.1% to 86.9% (Fig. 3e). From the plots visited in 2023, we found that pairs of the responsive parent PI 311785 and *inr-1/inr-1* NIL plants treated with OS derived from bean-fed fall armyworm demonstrated 40% reduction of attack to sentinel caterpillars by predatory *Polybia sp and Mischocyttaurus sp* wasps to *inr-1/inr-1* plants. Similarly in 2024, pairs of NILs treated with a physiologically relevant concentration or excess of In11 showed a 40% reduction of attack to *inr-1/inr-1* across two independent experiments, while there was no difference in visitation for mock treated plants (w + H_2_O) (Fig. 3f and g). These results demonstrate that INR mediates the In11-triggered recruitment of natural enemies in the field.

Taken together, our laboratory and field assays using the response to In11 in common bean landraces and near-isogenic lines (NILs) demonstrate that the defense responses upon In11 detection by the INR receptor mediate the interactions between plants, herbivorous pests, and predatory wasps in an agricultural ecosystem. In agreement with previous findings, we report that In11 is the primary elicitor of induced defenses found in herbivore OS, and that physiologically relevant concentrations of In11 elicit HIPV emission. The recruitment of predatory wasps to In11- or OS-treated plants in the field requires a functional INR receptor and is associated with the emission of specific volatile blends. Our findings confirm the ecological function of receptor-mediated HIPV emissions in legumes and represent further evidence that inducible plant volatiles can be harnessed for sustainable crop protection^40–43^.

## Materials and Methods

### Plant growth

Seeds from all common bean lines (*Phaseolus vulgaris*) were sown on Sunshine mix No. 4 and germinated in a reach-in PGC-FLEX growth chamber (Argus/Conviron) at 26°C, 70% relative humidity, and 500 μMol of light with a 12/12 h light/dark photoperiod. On day 13, plants with a fully expanded first trifoliate were transferred to a walk-in PGW40 chamber (Conviron) at 25°C, 50% relative humidity, and 500 µMol m^−2^ s^−1^ light with a 12/12 h light/dark photoperiod prior to experimentation.

### Genotyping and Resequencing

Samples were taken from primary leaves of common bean plants and flash frozen in liquid nitrogen. Frozen samples were then ground to a fine powder using a mixer mill (Retsch MM400, 2008, Part No. 20.745.0001) and genomic DNA extraction was performed using the NucleoSpin® Plant II kit (Machery-Nagel, 740770.520) according to the manufacturer’s instructions. The concentration and quality of the extracted DNA was assessed by NanoDrop One (ThermoFisher Scientific ND-ONE-W) and gel electrophoresis. Common bean accessions were genotyped using gene-specific primers (Extended data Table 2) flanking a ~1000 bp region that contained a 103 bp indel in the PvINR locus. PCR reactions were performed using gene-specific primers with DreamTaq PCR MasterMix (Thermo Fisher Cat. No. K1081). The PCR products were loaded onto a 1.5% agarose gel and run for 60 minutes at 120 V.

For resequencing, Illumina libraries were synthesized and 150 bp paired-end reads were sequenced with the NovaSeq 6000 instrument by Novogene Inc. Reads were mapped to the *Phaseolus vulgaris* reference genome using BWA-MEM (v 1.2.3) with default settings. Polymorphisms were called with FreeBayes (v 1.1)^44^ and relatedness calculated with TASSEL^45^

### Generation of near-isogenic lines (NILs)

Mature anthers from opened flowers of *Phaseolus vulgaris* W6 13807 containing the 103-base pair (bp) deletion in the *PvINR* locus (*inr-1*) were excised and used to pollinate the immature stigma from unpollinated, unopened host flowers of *Phaseolus vulgaris* PI 311785 without a deletion (*INR*). The resulting seeds were planted and genotyped as described. Heterozygous plants were kept and crossed with the recurrent line, PI 311785. This procedure was repeated until backcross six (BC6) lines were generated. For the BC five and six lines, heterozygous plants were allowed to self-pollinate and their offspring genotyped to select homozygous individuals with *INR/INR* or *inr-1/inr-1* genotypes.

### In11-induced ethylene gas production and gene expression on common bean

The In11 peptide (ICDINGVCVDA) was synthesized (GeneScript Inc) based on the cATP synthase sequence from cowpea (*Vigna unguiculata*) ^18^ and reconstituted in water. For ethylene gas measurements, the left and middle leaflet of the first trifoliate or the two primary leaves of a common bean plant were lightly wounded on both sides of the main vein using a fresh razor blade to remove the cuticle, and 10 μL of water alone or 1 μM In11 in 10 μL of water were distributed between the wounds using the tip of a pipette. After 1 hour, leaflets or primary leaves were excised at the petiole using a razor blade and placed in sealed tubes for 1 hour before sampling 1 mL of the headspace. Ethylene gas was measured as previously described^46^ with a gas chromatograph (HP 5890 series 2, supelco #13018-U, 80/100 Hayesep Q 3FT x 1/8IN x 2.1MM nickel) with flame ionization detection and quantified using a standard curve (Scott, 99.5% ethylene, Cat. No 25881-U).

For gene expression analyses, the middle leaflet of the first trifoliate or the primary leaves of a common bean plant were lightly wounded on both sides of the main vein using a fresh razor blade to remove the cuticle, and 10 μL of water alone or 1 μM In11 in 10 μL of water were distributed between the wounds using the tip of a pipette. After 1 hour, leaf samples were collected by removing the treated area with a 0.6 cm^2^ cork borer and flash frozen in liquid nitrogen. All samples were stored at −80°C until RNA extraction.

### In11-induced MAPK phosphorylation and immunoblotting

To measure In11-induced MAPK phosphorylation, we sprayed the first fully expanded trifoliate of 17-day-old common bean landraces and NILs with approximately 3 mL of a 0.01% Silwet L-77 (PhytoTech) solution with or without 1 μM In11. Leaf punches were collected from the three leaflets 15 minutes after treatment, alongside untreated controls, and combined before freezing in liquid nitrogen.

Total proteins were extracted using a 3X Laemmli (50 mM Tris-Cl pH 6.8, 6% SDS, 30% glycerol, 16% β-mercaptoethanol, and 0.006% bromophenol blue sample buffer) and separated by SDS-PAGE before transfer to a nitrocellulose membrane. To detect phosphorylated MAPK proteins, we used an anti phospho-p44/42 MAPK (Erk1/2) (Thr202/Tyr204) antibody (Cell signaling, #4370) at 1:2000 diluted in 1% Tris-buffered saline with Tween-20 (TBST) buffer, with secondary antibody anti-rabbit IgG-HRP conjugate (Sigma) at 1:10,000, followed by chemiluminescent visualization. Ponceau S solution (Sigma-Aldrich) staining was used to verify equal loading. The molecular weight of MAPK3 (42.66 kDa) and MAPK6 (45.06 kDa) were estimated based on their amino acid sequence.

### In11-induced volatile production

The seeds of the parental landrace and NILs were germinated on wet cotton in a Petri dish closed with Parafilm and covered with aluminum foil at 28°C for 48 hours. Subsequently, the seedlings were transplanted in individual cylindrical plastic pots filled with commercial soil (Einheitserdewerke Patzer Gebrüder Patzer GmbH & Co KG, Germany) and kept in a controlled growth chamber (L16:D8 light, 26°C 2, and 60% R.H.) until they were had two fully expanded trifoliates. The plants were watered every other day.

The three leaflets of a trifoliate in 12–16-day old plants were mechanically damaged by scratching the upper leaf surface with a razor blade and applying water (w + H_2_O) or 10 μL of 45 nM or 150 nM In11. The plants received the treatment twice: at 3:00 PM in the afternoon before the experiment and at 8:00 AM on the day of the experiment. The volatiles were collected as previously described^47^ 1 hour after the second treatment, for 3 hours. Briefly, the plants were individually placed in a glass bottle with clean and humidified air pushed inside through Teflon tubing at a rate of 0.8 L/min and the volatiles were trapped using 25mg of 80-100 mesh of HayeSep Q absorbent (Sigma, Switzerland) that was connected via Tygon tubing to a vacuum pump that pulled out the air at a rate of 0,5L/min. After 3 hours of volatiles collection each trapping filter was eluted with 100 μL of dichloromethane (Honeywell, Riedel-de Haën, DE) and 200 ng of n-nonyl acetate in 10 μL dichloromethane was added as internal standard. Finally, all the samples were stored in a −80°C freezer before analysis.

The samples were analyzed using a gas chromatograph (Agilent 7890B) coupled to a mass spectrometer (Agilent 5977B GC/MSD) in TIC mode. 2 μL of each sample was injected in pulsed spitless mode onto an Agilent HP-5MS column (30 m length x 0.25 mm diam. and 0.25m thick). After injection, the temperature program was the following: 40°C for 3min, increased to 100°C at a rate of 8°C/min and subsequently at 5°C/min until reaching 200°C and a post run of 3 min at 250°C. Helium was used as a carrier gas with a constant flow of 1.1 mL/min. For the identification of the volatiles, the mass spectrum of each compound was compared with the ones from two reference databases, NIST mass spectral library and PBM Quicksearch library (U.S. Department of Commerce). Identifications were further confirmed based on the spectra and retention times of authentic standards. The relative quantities (ng/h) of the most relevant compounds were calculated considering their response factors and internal standards peak areas.

### flg22-induced reactive oxygen species (ROS) accumulation

The flg22 peptide was synthesized (Genescript Inc) according to canonical flagellin sequence^48^ and reconstituted in water. For the assay, punches were taken from the primary leaves of common bean NILs with a 4 mm biopsy punch and floated overnight in 150 μL of H_2_O using individual cells of a white 96-well white bottom plate (BRANDplates F pureGrade S white). After overnight incubation, ROS production was measured upon addition of a 100 μL assay solution which contained 10 μg/mL luminol-horseradish peroxidase, 17 μg/mL luminol and H_2_O or flg22 to a final concentration of 2 μM. Luminescence was quantified with a TECAN SPARK plate reader every minute for 1 hour using an integration time of 500 ms, and the average ROS production for each plant was the mean of four leaf discs at each time point.

### Transcriptomics

To characterize transcriptional changes in response to wound and In11 treatment in common bean plants, we treated the middle leaflet of the first fully expanded trifoliate of *Phaseolus vulgaris* cv. Red Hawk by scratch wounding and adding water (w + H_2_O) or 1 μM In11 (w + In11). Samples were collected 1 hour later and flash frozen in liquid nitrogen. Total RNA extraction was performed using the Spectrum Plant RNA kit (Sigma) and mRNA was isolated with the NEBNext High Input Poly(A) mRNA Isolation Module (New England BioLabs). Paired-end Illumina 2×150 bp libraries were prepared with NEBNext Ultra™ RNA Library Prep Kit (New England BioLabs) and multiplexed with NEBNext® Multiplex Oligos for Illumina® (Index Primers Set 1). The sequencing was performed by Novogene Inc. For gene expression analyses, the reads were mapped to the reference *Phaseolus vulgaris* v2.1 genome available in Phytozome13^49^ and quantified using the –quantMode in STAR^50^. Differential gene expression analyses were performed using DESeq2^51^ implemented in R.

### RNA extraction, cDNA synthesis and qRT-PCR

Total RNA was extracted using the Trizol (Invitrogen) method. Briefly, frozen samples were ground to a fine powder using a mixer mill (Retsch MM400, 2008), and 900 μL of Trizol were added and homogenized using a vortex. The cell debris was separated by centrifugation, and the supernatant recovered in a clean tube. 200 μL of chloroform were added to the supernatant for phase separation. After centrifugation, the aqueous phase (top layer) was recovered, and the RNA was precipitated using an equal amount of isopropanol. RNA pellets were washed using 70% ethanol and resuspended in water. The concentration and quality of the extracted RNA was assessed by NanoDrop One (ThermoFisher Scientific ND-ONE-W) and gel electrophoresis, respectively. As DNA contamination was evident from the gel electrophoresis, the RNA was treated with TURBO DNA-free kit (Invitrogen) following manufacturer’s instructions. cDNA was synthesized using 1μg of RNA using the SuperScript IV Reverse Transcriptase Kit (Thermo Fisher, Cat. No. 18090050). qPCR reactions were performed using gene-specific primers with Power SYBR Green PCR Master Mix (Thermo Fisher, Cat. No. 4367659). Relative expressions were quantified using the ΔΔCq method, with ΔCq values normalized to the common bean UBQ gene and made relative to w + H_2_O treatment.

### Herbivory assays

First instar beet armyworm (*Spodoptera exigua*) larvae were obtained from Benzon Research Inc (Carlisle, PA). Upon receipt, larvae were incubated in their original packaging (with artificial diet) under dark conditions at 28°C for 36-48 hours to accelerate molting. 2^nd^ instar larvae were weighed using a digital scale (Mettler Toledo International Inc) and placed in plastic cups without artificial diet. Once all larval masses were recorded, one single larva was placed on the first trifoliate or primary leaf of *INR/INR* and *inr-1/inr-1* plants, and leaves were enclosed in a transparent mesh bag to contain the larva. Infested plants were kept in a growth chamber at 25°C, 50% RH, and 500 µMol m^−2^ s^−1^ light, and larvae weights were recorded again 5 days later.

### Sentinel-prey field assay

Field assays were conducted in 2023 and 2024 in a 925 m^2^ agricultural field near Bajos de Chila, Oaxaca, Mexico (15,91818° N, 97,16401° W) (Fig. 3).

For the 2023 field season, seeds of the parental PI 311785 landrace and the *inr-1/inr-1* NIL were sown 1.5 L plastic pots containing field-collected soil. To prevent herbivory, plants were grown in large mesh cages (2 × 2 × 4 m; BioQuip, CA, USA) for 13 days and watered every other day. Plants were treated by removing 2 cm^2^ of cuticle on the adaxial side of an expanded leaflet with sandpaper and adding water (w + H_2_O) or fresh oral secretions (w + OS) derived from bean-fed fall armyworm larvae (*Spodoptera frugiperda*). The treatments were imposed the evening before and the morning of the experiment. After the second damage treatment, one randomly selected freeze-killed caterpillar (defrosted) was pinned to the center of each leaf (two caterpillars per plant). At 9:00 am, pairs of potted PI 311785 and *inr-1/inr-1* treated by either wounding and adding water (w + H_2_O) or OS (w + OS) and with pinned caterpillars were arranged in six spatial blocks in plots within the experimental field. Paired pots were approximately 30 cm apart.

Blocks were visited continuously over 90 min, and the order in which sentinel caterpillars from the different treatments were subject to committed visits by one or more wasps in each block was recorded. The experiment was repeated a second day using the same plants but arranged in different treatment pairs.

For the 2024 field season, seeds of *INR/INR* and *inr-1/inr-1* sibling NILs were grown as described above. 13-day old plants were treated at 7:30 am by removing 2 cm^2^ of cuticle on the adaxial side of two opposing leaflets on the 1^st^ fully expanded trifoliate with a razor blade and adding 45 nM In11 (w + In11) or water (w + H_2_O). After imposing the same treatment 24 hours later, freeze-killed fall armyworm caterpillars were pinned to the treated leaflets as described above.

Beginning at approximately 08:00 am, treatment pairs (*w* + In11 _*INR/INR*_ and *w* + In11 _*inr-1/inr-1*_) were observed for wasp visitation in five spatial blocks of the experimental field. For each pair, we recorded the time of the first wasp visit, defined as a committed attack on a larva, as well as the species of the first wasp to arrive. Following the initial visit in each block, we recorded the number of wasps on each plant at 5, 15, 30, 45, 60, 75, and 90 minutes. The experiment was repeated on a second day using the same plants, but with treatments reassigned to new pairings. Three days later, the trial was repeated on a new set of In11-treated plants, and seven days after that with plants that were only water treated after the razorblade-inflicted damage.

### Statistical analysis

All statistical analyses were conducted in R (R version 4.3.2 or 4.3.3, R Core Team 2023). We first confirmed that the residuals fulfilled assumption of normality using a Shapiro Wilk Test and qq plots and then implemented t-test to detect differences in the mean ethylene gas accumulation, gene expression, and mean growth rate of beet armyworm caterpillars feeding on *inr-1/inr-1* or *INR/INR* NILs. Generalized linear models were used to assess the effect of treatments on volatile emissions. A Gamma error distribution was used to account for the non-normality of emission data. Estimated marginal means were calculated using the emmeans package (Lenth, 2023), and pairwise comparisons were conducted with a false discovery rate (FDR) correction for multiple testing.

To visualize overall differences in volatile profiles across treatments, we conducted principal component analysis (PCA). Volatile compound emissions were log-transformed and scaled to unit variance prior to analysis.

Generalized linear mixed models using the glmmTMB package^52^ were used to assess the effect of treatment (2023, OS; 2024, In11 and H_2_O) on the removal of sentinel caterpillars from bean plant pairs in the field. To account for the hierarchical structure of the experiment, we included random effects for plots (plant pairs) nested within spatial blocks, which were in turn nested within each trial. Date was included as a fixed effect in the model, as only two trial days were used and random effects with fewer than five levels are generally discouraged due to unstable variance estimates^53^.

## Data availability

All re-sequencing and transcriptomics data is available at the National Center for Biotechnology Information (NCBI) under PRJNA1278804 and PRJNA1272629, respectively. All other data are available in the manuscript or the supplementary materials.

## Acknowledgments

We thank members of the Nemhauser, Imaizumi and Di Stilio labs for conversation and feedback, and Elly Poretsky and Yezhang Ding for manuscript comments. We thank Jim Kelly for providing *Phaseolus vulgaris* cv. red hawk seeds.

## Author contributions

N.G.P. and P.G-T contributed equally to this work. A.D.S, T.C.J.T, B.B, N.G.P and P.G-T conceived the project. A.F.C and B.B (UW) generated NILs. N.G.P characterized the NILs in the laboratory. C.M.A and P.G-T collected and characterized the volatiles. D.W performed the MAPK phosphorylation assay. A.D.S, B.B (UW), T.C.J.T, P.G-T and B.B recorded wasp visitation in the field. N.G.P and P.G-T analyzed the data and generated the figures. N.G.P and A.D.S wrote the manuscript with contributions from all authors. All authors read and edited the manuscript.

## Extended data

**Extended data Figure 1.**
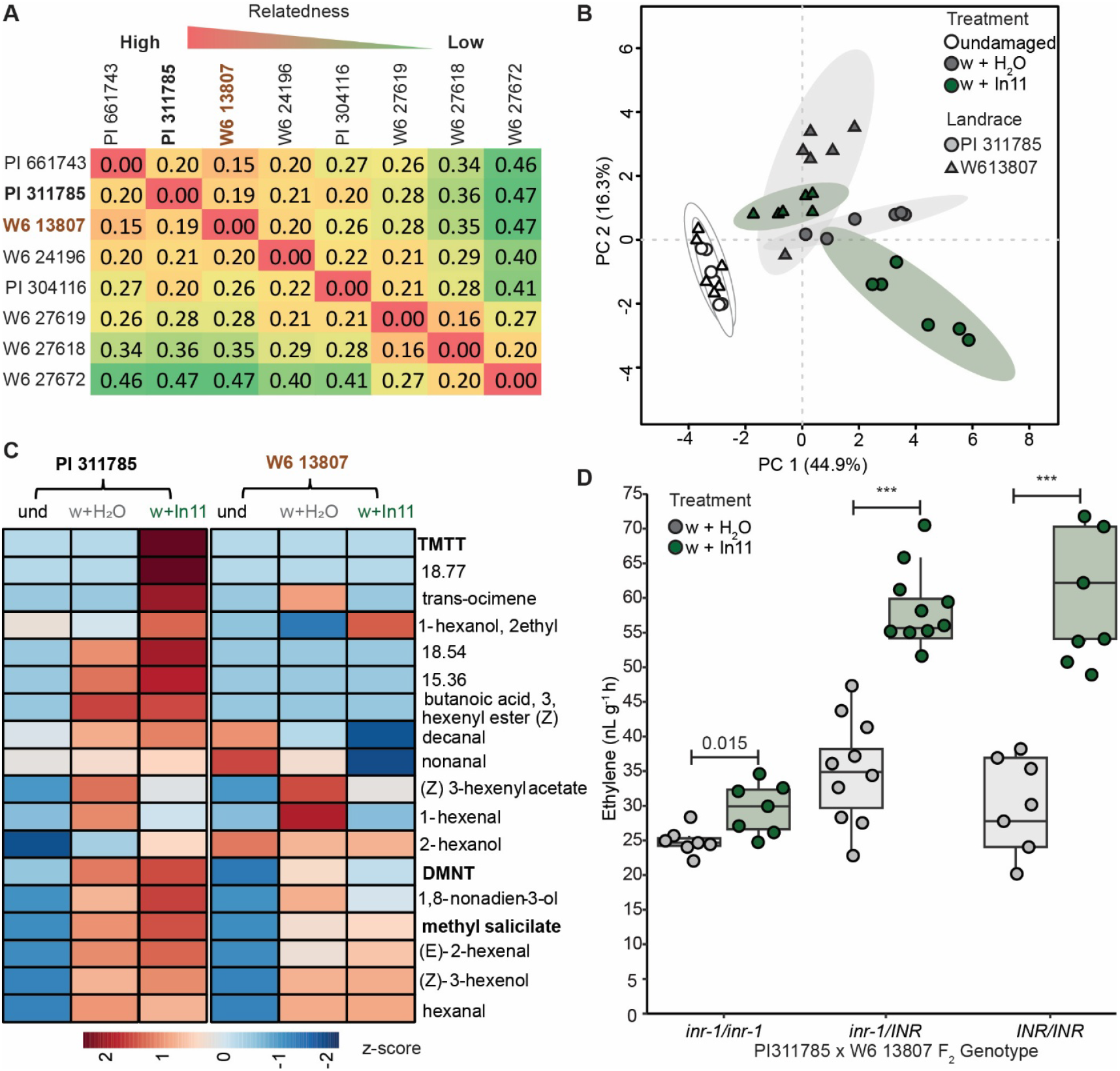
(**A)** Relatedness matrix of Honduran common bean landraces used in the ethylene screen. Cross-compatible In11-responsive (PI 311785) and non-responsive (W6 13807) landraces are indicated in bold. (**B)** Principal component analysis (PCA) of the volatile profiles from undamaged and 2 hours after wounding (w + H_2_O) or In11 (wound + In11) treatment in selected common bean landraces (n=6). (**C)** Relative abundance of specific compounds in the volatile blends. **D**. Ethylene production of F2 individuals from a PI 311785 x W6 13807 cross.

**Extended data Figure 2.**
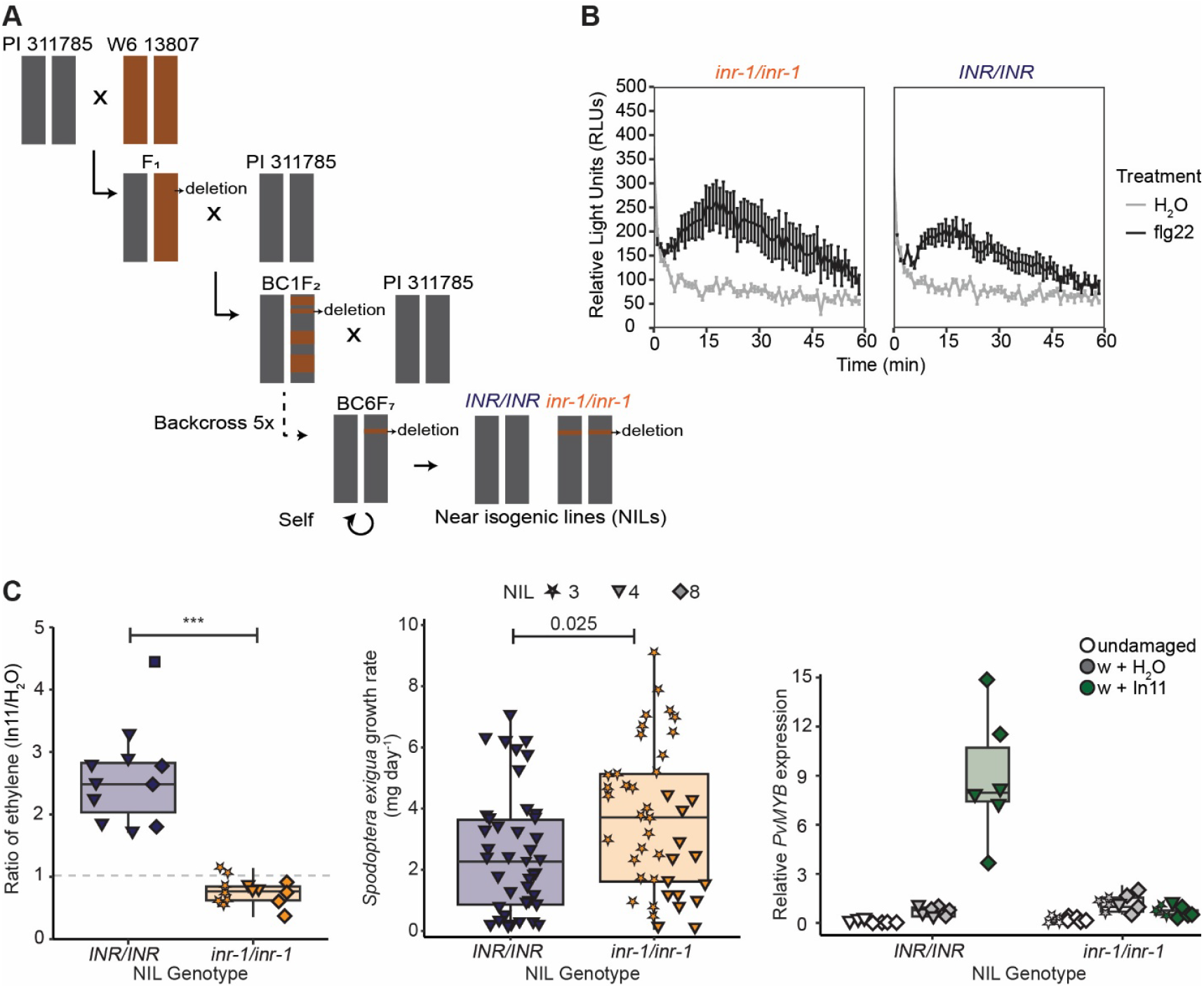
flg22-induced reactive oxygen species (ROS) burst in common bean NILs. **(A)** Back-crossing scheme of sexually compatible In11-responsive (PI 311785) and non-responsive (W6 13807) landraces to generate *INR/INR* and *inr-1/inr-1* near-isogenic lines (NILs) in the PI 311785 background. NILs labeled as lines 3, 4, and 7 derived from three independent initial crosses are used in the work, each unresponsive to In11. (**B)** Oxidative burst produced by primary leaves from common bean NIL 4 treated with 2 µM flg22 peptide and measured in relative luminescence units (RLU). Results shown are means ± s.d. (n = 3 plants)

**Extended data Figure 3.**
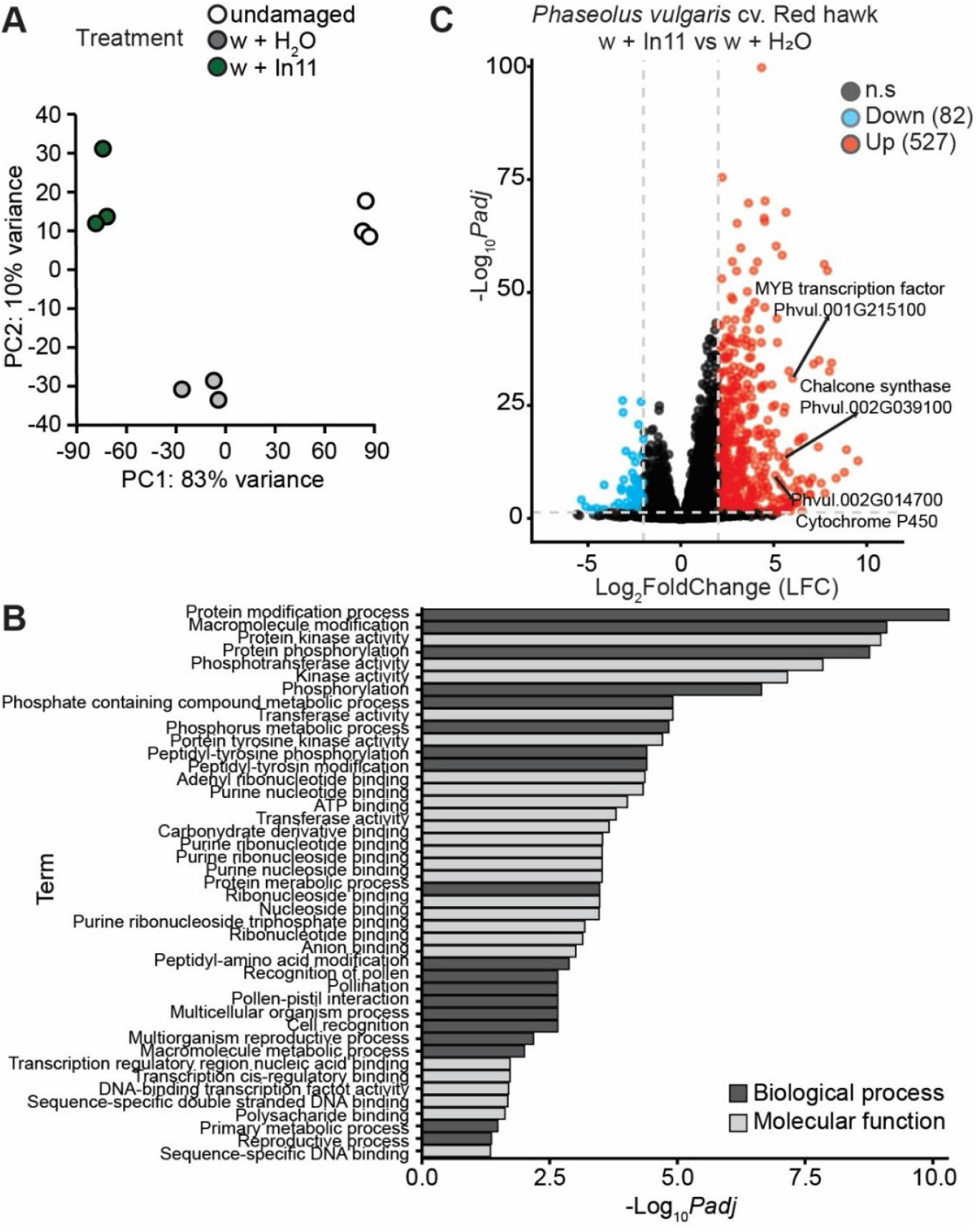
RNAseq analysis of In11-treated common bean plants. (**A)** Principal component (PC) analysis of differentially expressed genes (DEGs) across all samples. (**B)** Volcano plot of DEGs 1 hour after wounding (w) + 1 µM In11 vs wounding alone (w + H2O) treatment in *Phaseolus vulgaris* cv. Red Hawk. Selected marker genes *PvMYB, PvCHS*, and *PvCYP* are indicated. (**C)** Enriched Gene Ontology (GO) categories among In11-upregulated genes.

**Extended data Table 1.** Ethylene response ratios of *Phaseolus vulgaris* plants scratch wounded and treated with wounding (w) + H_2_O or w + In11.

| Variety | w + In11 / w + H<sub>2</sub>O |
| --- | --- |
| W6 13807 | 0.7051507095 |
| W6 17491 | 0.7453989813 |
| PI 638866 | 1.243128964 |
| pi 309831 | 1.288296976 |
| PI 550055 | 1.316718588 |
| PI 570661 | 1.356811961 |
| W6 21064 | 1.38740458 |
| PI 430199 | 1.454239507 |
| PI 476742 | 1.499436253 |
| PI 209493 | 1.516271532 |
| PI 432617 | 1.642282661 |
| PI 313536 | 1.673658614 |
| PI 535272 | 1.679866869 |
| PI 318694 | 1.736723542 |
| PI 325669 | 1.789803615 |
| PI 313293 | 1.83336787 |
| PI 632862 | 1.914207886 |
| PI 638873 | 1.98050087 |
| PI 413673 | 1.985270114 |
| PI 313502 | 2.019751588 |
| W6 27747 | 2.099041534 |
| w6 18811 | 2.116945814 |
| W6 27621 | 2.226574136 |
| PI 163120 | 2.376400387 |
| PI 310511 | 2.492743214 |
| PI 642111 | 2.731270054 |
| PI 395776 | 2.739277503 |
| PI 186492 | 2.740451047 |
| PI 313462 | 2.777926358 |
| PI 196299 | 2.781456729 |
| PI 583354 | 2.871189599 |
| PI 310573 | 2.910958904 |
| PI 476693 | 2.940380315 |
| PI 200963 | 2.955424685 |
| PI 638869 | 3.025578812 |
| PI 53713 | 3.114577362 |
| W6 13744 | 3.209294302 |
| W6 27749 | 3.301886792 |
| PI 550027 | 3.331991952 |
| PI 310524 | 3.409215866 |
| PI 325672 | 3.417773238 |
| PI 309696 | 3.452404514 |
| PI 263598 | 3.588614024 |
| W6 27750 | 3.599863004 |
| PI 638853 | 3.70171144 |
| PI 209493 | 3.768890894 |
| PI 313340 | 3.774676749 |
| PI 325689 | 3.849158957 |
| PI 200967 | 3.959417388 |
| PI 352750 | 4.060400933 |
| PI 313316 | 4.066894561 |
| PI 304116 | 4.097433655 |
| PI 621064 | 4.282001477 |
| pi 304112 | 4.2989178 |
| W6 27753 | 4.325791855 |
| PI 313315 | 4.401146784 |
| PI 441831 | 4.455930053 |
| PI 1325672 | 4.507188498 |
| pi 661818 | 4.510982792 |
| PI 262893 | 4.66915814 |
| Red Hawk | 4.683830317 |
| PI 535363 | 4.7744713 |
| PI 313446 | 4.801101467 |
| PI 592898 | 4.889059597 |
| PI 308893 | 4.948415259 |
| PI 430213 | 5.135408788 |
| PI 262893 | 5.138841672 |
| PI 325678 | 5.429767255 |
| PI 1313498 | 5.61631427 |
| PI 406789 | 5.746853999 |
| PI 406788 | 5.824502621 |
| PI 577682 | 5.85562933 |
| PI 561473 | 5.979487179 |
| PI 196466 | 6.089983467 |
| PI 196466 | 6.090614887 |
| PI 313380 | 6.437524691 |
| W6 35979 | 6.497070652 |
| PI 313498 | 7.454761905 |
| PI 1549860 | 7.522222222 |
| PI 151615 | 8.167234948 |
| PI 549870 | 8.245726496 |
| PI 638835 | 8.524903715 |
| PI 549876 | 8.953109969 |
| PI 310911 | 9.854140915 |
| Black Valentine | 10.22581817 |
| pi 441834 | 10.86619027 |
| PI 310891 | 14.74923547 |
| PI 549860 | 15.13686667 |
| PI 470938 | 15.22044728 |
| PI 638903 | 16.48559671 |
| PI 352745 | 16.83826026 |

**Extended data Table 2.**
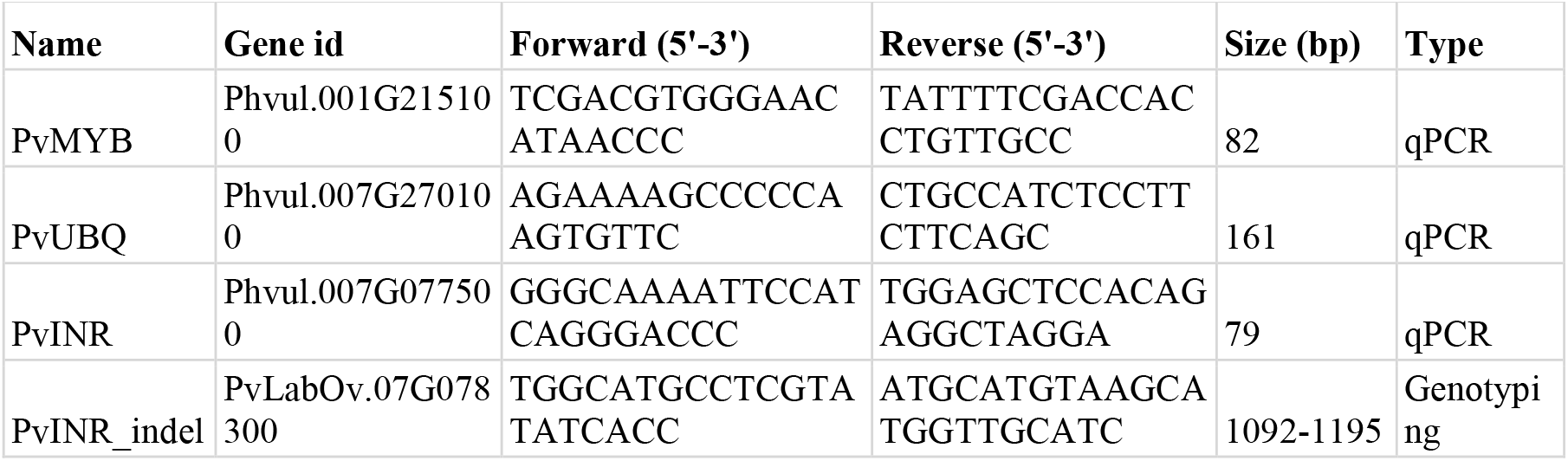
List of primers used in this study.

